# The cohesin ring uses its hinge to organize DNA using non-topological as well as topological mechanisms

**DOI:** 10.1101/197848

**Authors:** Madhusudhan Srinivasan, Johanna C. Scheinost, Naomi J. Petela, Thomas G. Gligoris, Maria Wissler, Sugako Ogushi, James Collier, Menelaos Voulgaris, Alexander Kurze, Kok-Lung Chan, Bin Hu, Vincenzo Costanzo, Kim A. Nasmyth

## Abstract

As predicted by the notion that sister chromatid cohesion is mediated by entrapment of sister DNAs inside cohesin rings, there is a perfect correlation between co-entrapment of circular minichromosomes and sister chromatid cohesion in a large variety of mutants. In most cells where cohesin loads onto chromosomes but fails to form cohesion, loading is accompanied by entrapment of individual DNAs. However, cohesin with a hinge domain whose positively charged lumen has been neutralized not only loads onto and translocates along chromatin but also organizes it into chromatid-like threads, despite largely failing to entrap DNAs inside its ring. Thus, cohesin engages chromatin in a non-topological as well as a topological manner. Our finding that hinge mutations, but not fusions between Smc and kleisin subunits, abolish entrapment suggests that DNAs may enter cohesin rings through hinge opening. Lastly, mutation of three highly conserved lysine residues inside the Smc1 moiety of Smc1/3 hinges abolishes all loading without affecting cohesin’s initial recruitment to *CEN* loading sites or its ability to hydrolyze ATP. We suggest that loading and translocation are mediated by conformational changes in cohesin’s hinge driven by cycles of ATP hydrolysis.

## Introduction

Smc/kleisin complexes facilitate chromosome segregation in bacteria (Burmann and Gruber, 2015) as well as eukaryotes (Schleiffer et al., 2003). Eukaryotes possess three types: condensin, which mediates chromatid formation during mitosis (Hirano et al., 1997), cohesin, which confers cohesion between sisters (Guacci et al., 1997; Michaelis et al., 1997), and the less well understood Smc5/6 complex (Haering and Gruber, 2016). Though initially identified as being essential for holding the sister chromatids together, cohesin shares with condensin the ability to organize DNA into chromatids, albeit during interphase, with loops emanating from an Smc/kleisin axis (Rhodes et al., 2017; Tedeschi et al., 2013). It has been suggested that cohesin and condensin organizes DNAs into chromatids by capturing small loops of DNA and then extruding them in a processive manner (Nasmyth, 2001). This concept, now known as loop extrusion (LE), has recently been embellished to explain the pattern of intra-chromosomal interactions during interphase, as measured by Hi-C, and the process by which interactions between enhancers and distant promoters are regulated by the insulation factor CTCF (Fudenberg et al., 2016; Sanborn et al., 2015). Because LE is thought also to apply to chromosome segregation in bacteria (Wang et al., 2017), to formation of bivalent chromosomes during meiosis, and to X chromosome dosage compensation in worms, it is of great importance to understand how Smc/kleisin complexes organize DNA topology.

At cohesin’s core is a heterotrimeric ring containing a pair of SMC proteins, Smc1 and Smc3, and an α-kleisin subunit Scc1. Smc1 and Smc3 are rod shaped proteins containing 50 nm long intra-molecular anti-parallel coiled coils with a hinge/dimerization domain at one end and an ABC-like ATPase head domain composed of the protein’s N- and C-terminal sequences at the other. They bind each other via their hinges to form V-shaped heterodimers whose apical ATPases are interconnected by a single Scc1 polypeptide (Gruber et al., 2003; Haering et al., 2002). Scc1’s N-terminal domain (NTD) forms a four helical bundle with the coiled coil emerging from Smc3’s ATPase (Gligoris et al., 2014) while its winged helical C-terminal domain (CTD) binds to the base of Smc1’s ATPase (Haering et al., 2004). Bacterial Smc/kleisin complexes form similar structures, suggesting that asymmetric ring formation is a universal feature (Burmann et al., 2013).

Together with the finding that cleavage of Scc1 by separase triggers cohesin’s dissociation from chromosomes and sister chromatid disjunction at the onset of anaphase (Uhlmann et al., 1999), the discovery that cohesin forms a ring raised the possibility that it holds DNAs together using a topological principle (Haering et al., 2002; Nasmyth, 2001). According to this notion, entrapment of individual DNAs inside cohesin rings constitutes the mechanism by which cohesin associates with chromosomes while co-entrapment of sister DNAs mediates sister chromatid cohesion. This is known as the ring model.

The finding that cohesin’s association with circular DNA is destroyed by restriction endonuclease cleavage as well as by endo-proteolytic cleavage of Scc1 (Ivanov and Nasmyth, 2005; Murayama and Uhlmann, 2014) is consistent with the above model. However, it neither definitively proves topological entrapment nor addresses the nature of cohesion between sister DNAs. These two issues have only become amenable due to elucidation of the molecular structures of the Smc1/Smc3, Smc3/Scc1, and Scc1/Smc1 interfaces, which permitted insertion of cysteine pairs that can be chemically crosslinked using bi-functional thiol-specific reagents (Haering et al., 2008). Crosslinking of all three interfaces creates circular Smc1-Smc3-Scc1 polypeptides within which either individual or indeed pairs of circular DNAs would be trapped in a manner resistant to SDS treatment. The chemical catenation of individual DNAs in this manner causes them to migrate slower during gel electrophoresis (catenated monomers or CMs) while catenation of sister DNAs causes them to migrate as dimers (catenated dimers or CDs), even if the two DNAs are not otherwise intertwined (Gligoris et al., 2014).

According to the ring model, cohesin’s association with and dissociation from chromosomes involves entry and exit of DNAs into and from rings, respectively. Cohesin dissociates from chromosomes after cleavage of Scc1 by separase at the onset of anaphase (Uhlmann et al., 1999) but at other stages of the cycle in a manner involving dissociation of the Smc3/Scc1 interface (Beckouet et al., 2016). This separase-independent releasing activity depends on engagement of Smc1 and Smc3 ATPase heads (Elbatsh et al., 2016), binding to Scc1 of a pair of hookshaped Hawk proteins (Wells et al., 2017), namely Scc3 (Roig et al., 2014) and Pds5 (Lee et al., 2016), both of which bind Wapl, a third releasing activity subunit (Kueng et al., 2006). Release also depends on a pair of highly conserved lysine residues (K112 and K113) on Smc3’s ATPase, whose modification by the acetyl transferase Eco1 during S phase abolishes release (Beckouet et al., 2016; Unal et al., 2008), thereby helping to maintain cohesion until the onset of anaphase. It is currently presumed that transient dissociation of the Smc3/Scc1 interface mediates release by creating a gate through which previously entrapped DNAs exit the cohesin ring.

The mechanism by which cohesin loads onto chromosomes is even less well understood. It too requires engagement of Smc1 and Smc3 ATPase heads as well as subsequent ATP hydrolysis (Arumugam et al., 2003; Weitzer et al., 2003). Neither Pds5 nor Wapl are necessary (Chan et al., 2013; Petela et al., 2017) but instead a separate complex containing Scc4 bound to the NTD of another Hawk protein called Scc2 is essential (Chao et al., 2015; Ciosk et al., 2000; Hinshaw et al., 2015), as might be Scc3 (Toth et al., 1999). Important insight into the process stemmed from discovery that the point centromeres of *S. cerevisiae* (CENs) which contain only a single CENP-A containing nucleosome, direct loading of most chromosomal cohesin within a surrounding 60 kb window (peri-centric cohesin) (Hu et al., 2011; Weber et al., 2004).

How DNA enters the cohesin ring during a loading reaction is controversial. Neither fusion of Smc3’s ATPase to Scc1’s NTD nor fusion of Smc?s ATPase to Scc1’s CTD is lethal, suggesting that entry through a unique entry gate created by dissociation of one or the other Smc/kleisin interface is not obligatory for loading (Gruber et al., 2006). The suggestion that DNAs enter via the Smc3/Scc1 interface under the influence of Wapl and Pds5 (Murayama and Uhlmann, 2015) is hard to reconcile with the above findings, as is the fact that neither Pds5 nor Wapl are necessary for loading in budding yeast (Chan et al., 2013), fission yeast (Bernard et al., 2008), plants (De et al., 2014) or animal cells (Tedeschi et al., 2013). Whether or not it is the entry gate, the hinge clearly has a crucial role during the loading reaction because complexes whose hinges contain a mutation that weakens but does not eliminate dimerization in vivo (Smc1F584R) or whose hinges have been replaced by a different dimerization domain cannot undergo even the first step in the loading reaction, namely association of hydrolysis-defective complexes with CENP-A loading sites (Hu et al., 2011). Furthermore, constraining the Smc1 and Smc3 hinge domains compromises loading and the establishment of sister chromatid cohesion (Gruber et al., 2006).

Two key questions are therefore outstanding. Is cohesion really mediated by coentrapment of DNAs inside cohesin rings and is loading per se mediated by entrapment of individual DNAs? If the answer to either is affirmative, then cohesin must have an entry gate and its identification is of paramount importance. By analyzing cells at different stages of the cell cycle or with a variety of mutations, we show that co-entrapment of sister DNAs within cohesin rings invariably accompanies sister chromatid cohesion. On the contrary, although entrapment of individual DNAs normally accompanies loading, we describe a situation where this does not apply, namely a cohesin mutant with a hinge whose positively charged lumen has been neutralized by five mutations (*smc1DD smc3AAA*). The anomalous behavior of this hinge mutant proves that cohesin is able to load onto and translocate along chromosomes without associating with them in a topological manner. It also confirms that topological entrapment is essential for sister chromatid cohesion. Remarkably, *smc1DD smc3AAA* cohesin can still fold DNAs into chromatid-like threads, suggesting that it may be capable of LE and that this process does not require strict topological entanglement of cohesin with DNA.

Equally remarkable, we describe a triple mutation that replaces by aspartic acid three highly conserved lysines from Smc1 within the hinge’s lumen that greatly reduces cohesin’s association with chromosomes despite associating with Scc2 at *CEN* loading sites and being fully active as an ATPase. The behavior of this *smc1DDD* mutation implies that changes in the conformation of cohesin’s hinge that normally accompany ATP hydrolysis are essential for completion of the loading reaction. We suggest that both topological and non-topological modes of cohesin loading/translocation depend on changes in cohesin’s Smc1/3 hinge domain that respond to changes in the state of its ATPase domains.

## Results

### Entrapment of sister DNA molecules by hetero-trimeric cohesin rings

To measure DNA entrapment by cohesin, we created a pair of strains containing 2.3 kb circular minichromosomes: a 6C strain with cysteine pairs at all three ring subunit interfaces (Smc1G22C K639C, Smc3E570C S1043C, Scc1A547C C56) and a 5C strain lacking just one of these (Scc1A547C). Exponentially growing cells were treated with BMOE, which circularizes 20-25% of 6C cohesin rings (Gligoris et al., 2014), and DNAs associated with cohesin immunoprecipitates separated by agarose gel electrophoresis following SDS denaturation. Southern blotting revealed two forms of DNA unique to 6C cells: one that migrates slower than monomeric supercoiled DNAs (CMs) and a second that migrates slower than DNA-DNA concatemers (CDs) (Fig. 1AB). Little if any minichromosome DNA is detected in cells lacking the affinity tag on cohesin (Fig. 1B, S1A). Importantly, electrophoresis in a second dimension following proteinase K treatment confirmed that both forms consist of monomeric supercoiled DNAs. CMs are single DNA molecules trapped within cohesin rings while CDs contain a pair of sister DNAs trapped within tripartite cohesin rings (Fig. S1B).

**Figure 1.**
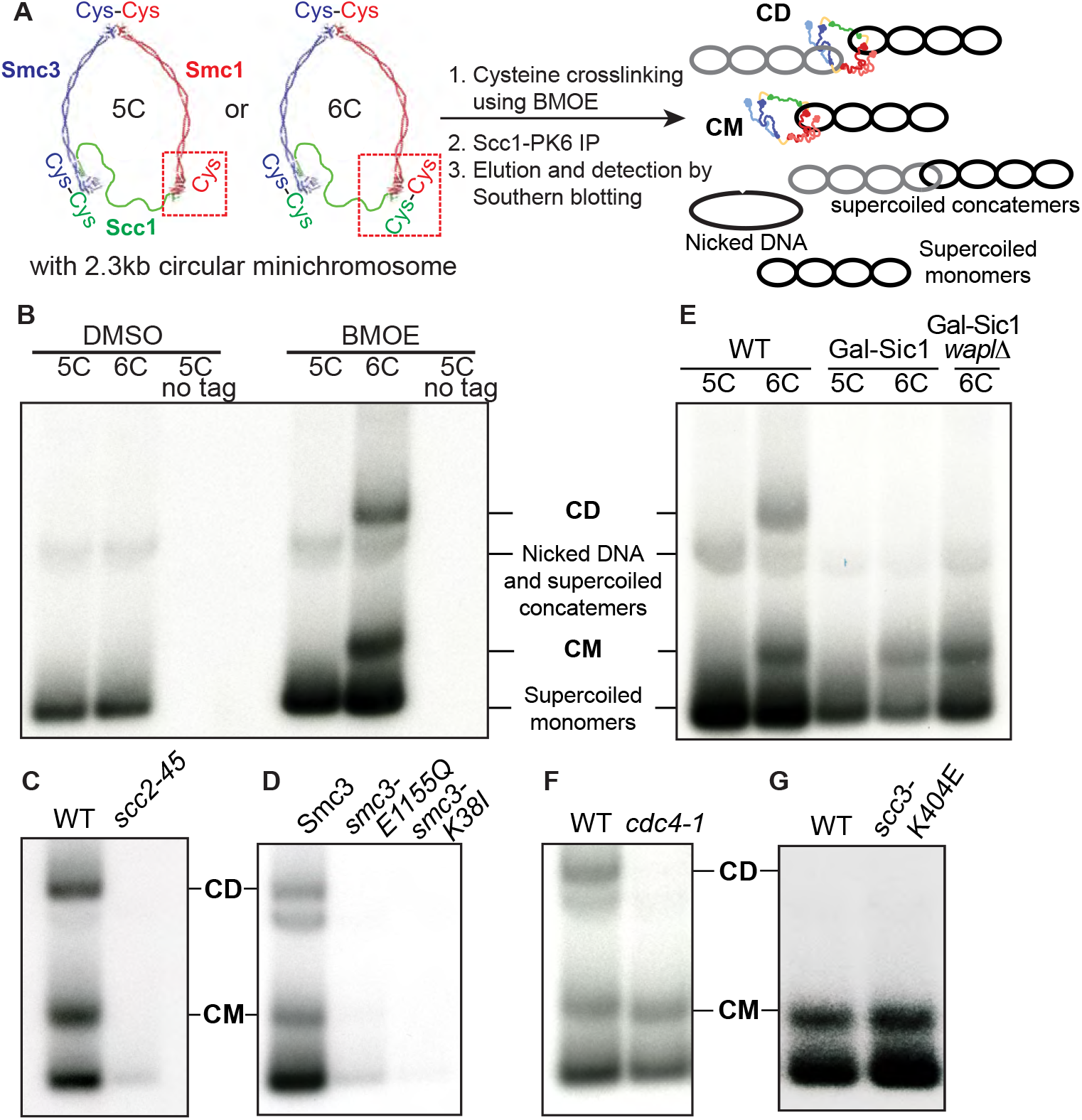
Entrapment of single and sister DNA molecules by hetero-trimeric cohesin rings. **(A)** Schematic of the minichromosome IP assay to measure DNA entrapment by cohesin. 6C strains with cysteine pairs at all three ring subunit interfaces (2C Smc3: E570C S1043C, 2C Smc1: G22C K639C and 2C Scc1 C56 A547C) and 5C strains lacking just one of these cysteines (Scc1 A547C) and carrying a 2.3 kb circular minichromosome were subjected to in vivo crosslinking with BMOE. DNAs associated with cohesin immunoprecipitates were denatured with SDS and separated by agarose gel electrophoresis. Southern blotting reveals supercoiled monomers, nicked and supercoiled concatemers of the minichromosome along with two forms of DNA unique to 6C cells, termed CMs and CDs. **(B)** Exponentially growing strains K23644 (5C), K23889 (6C) and K23890 (5C, no cohesin tag) were subjected to the minichromosome IP assay as described in (A). The positions of the different DNA species are marked in the Southern blot. See also Figure S1A and S1B. **(C)** Wild type (K23889) and *scc2-45* (K24267) 6C strains were arrested in G1 with α-factor at 25°C in YPD medium and released into medium containing nocodazole at 37°C to arrest cells in G2, and then subjected to minichromosome IP. See also Figure S1C. **(D)** Exponentially growing 6C strains containing ectopically expressed versions of 2C Smc3: K24173 (wild type Smc3), K24174 (smc3 E1155Q) and K24175 (smc3 K38I) were subjected to minichromosome IP assay. See also Figure S1D. **(E)** Strains K23644 (5C), K23889 (6C), and those expressing galactose-inducible nondegradable sic1 K23971 (5C), K23972 (6C) and K23451 (6C *wplΔ*) were arrested in late G1 as described in STAR methods and subjected to minichromosome IP assay. See also Figure S1E. **(F)** Wild type (K23889) and *cdc4-1* (K24087) 6C strains were arrested in G1 at 25°C in YPD medium and released into YPD medium containing nocodazole at 37°C to arrest cells in G2, and subjected to minichromosome IP assay. See also Figure S1F. **(G)** Non-cleavable 2C scc1 was expressed in G1 arrested wild type (K24695) and *scc3 K404E* mutant (K24697) strains, followed by minichromosome IP assay. See also Figure S1G.

To address whether CMs and CDs correspond to loading and cohesion, respectively, we measured CM and CD formation in a variety of mutants and cell cycle states. We first asked whether mutants defective in loading fail to create CMs and CDs. Accordingly, *scc2-45* cells failed to co-precipitate minichromosomes after undergoing DNA replication in the absence of functional Scc2 loader (Fig. 1C). Likewise, a version of 6C cohesin that can bind but not hydrolyze ATP (Smc3E1155Q) and a version that cannot even bind ATP (Smc3K38I) failed to co-precipitate minichromosomes (Fig. 1D). It is important to note that Smc3E1155Q cohesin associates with centromeres with a very high efficiency, whether measured by live imaging (Hu et al., 2011) or by calibrated ChiP-seq (Hu et al., 2015), and yet it largely fails to immuno-precipitate minichromosome DNA, which correlates with its failure to associate with chromatin in a fashion sufficiently stable to survive minichromosome isolation.

### Cohesin entraps individual DNAs before DNA replication

If cohesion were mediated by co-entrapment of sister DNAs within cohesin rings, then CDs should be detected only in cells that have undergone DNA replication. Likewise, if cohesin loading involved entrapment of individual DNAs, then CMs should be detected in cells that load cohesin onto chromosome prior to replication. Expression of a non-degradable version of the Cdk1 inhibitor Sic1 or inactivation of the F-box protein Cdc4, which is necessary for Sic1 degradation, prevents cells from entering S phase but allows inactivation of separase and the burst of cohesin loading that follows activation of *SCC1* gene expression in late G1. In both cases, the failure to degrade Sic1 (following release from an α-factor induced G1 arrest) was accompanied by CM but not CD formation (Fig. 1EF). Expression of a version of Scc1A547C C56 that cannot be cleaved by separase in α-factor arrested G1 cells also led to formation of CMs but not CDs (Fig. 1G, left lane). CM accumulation was marginally greater in cells whose releasing activity is compromised by a mutation *(scc3K404E)* that abolishes the interaction between Wapl and Scc3 (Beckouet et al., 2016) (Fig. 1G, right lane). These results demonstrate that CMs can form in the absence of CDs and that they are not merely a byproduct of CD formation.

### Sister chromatid cohesion is generated by entrapment of sister DNAs within individual cohesin rings

CMs but not CDs formed with high efficiency when ts *eco1-1* (Fig. 2A) mutant cells underwent DNA replication at 37°C. Loading is known to take place with high efficiency in such cells despite their failure to create stable cohesion (Chan et al., 2013). The lack of CDs suggests that entrapment of sister DNAs inside cohesin rings may be an intrinsic aspect of sister chromatid cohesion. If so, inactivation of releasing activity, which suppresses the lethality of *eco1-1* mutants and at least partly restores establishment of cohesion should also restore CD formation. This is indeed the case. *wpl1Δ*, which permits proliferation of *eco1-1* cells at 37°C, also restored efficient CD formation (Fig. 2A).

**Figure 2.**
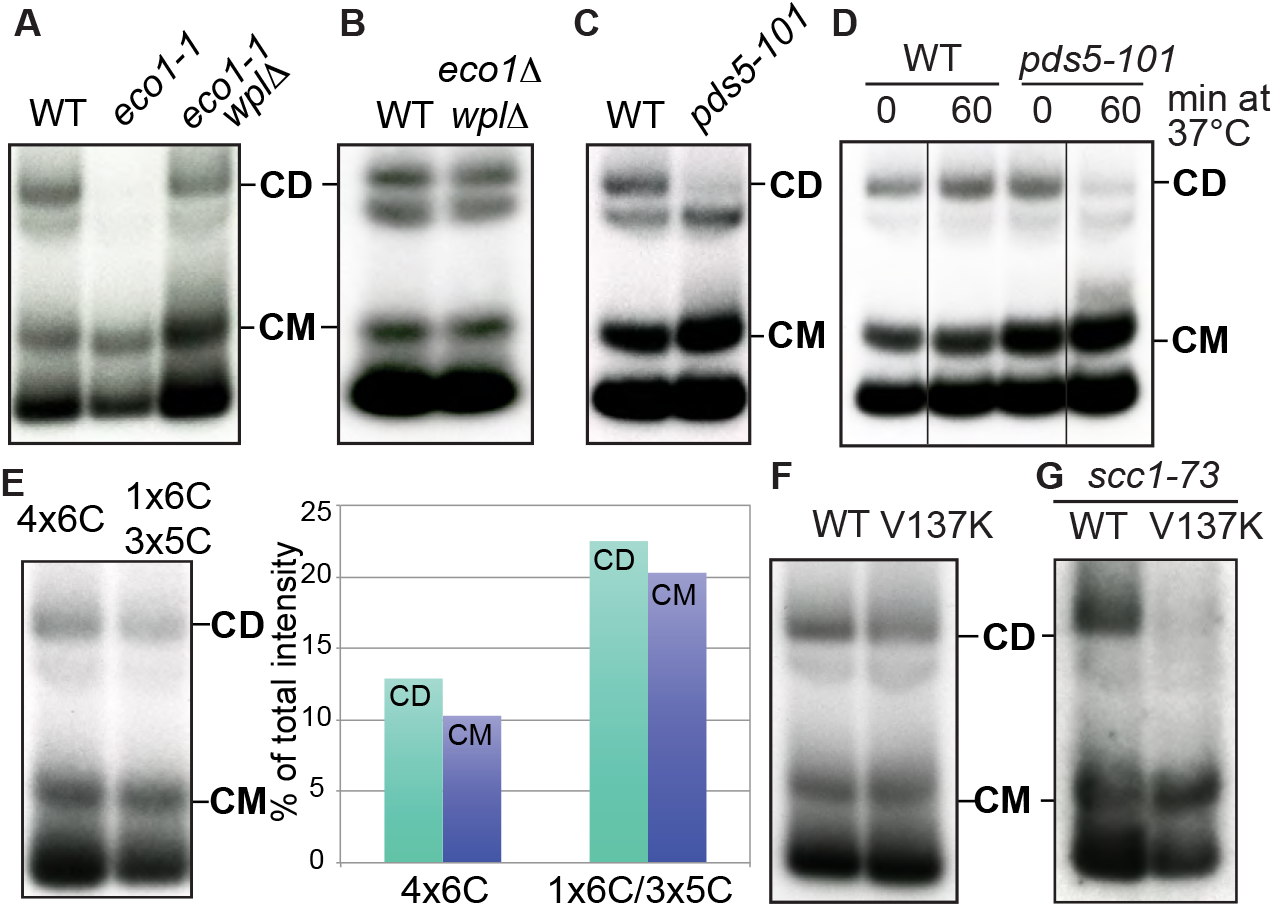
Sister chromatid cohesion is generated by entrapment of sister DNAs within individual cohesin rings. **(A)** Wild type (K23889), *eco1-1* (K23579) and *eco1-1 wplΔ* (K23578) 6C strains were arrested in G1 at 25°C in YPD medium, released into YPD medium containing nocodazole at 37°C to arrest cells in G2 and subjected to minichromosome IP assay. **(B)** Minichromosome IP assay of exponentially growing wild type (K23889) and *eco1Δ wplΔ* (K25287) 6C strains. **(C)** Wild type (K23889) and *pds5-101* (K24030) 6C strains were arrested in G1 at 25°C in YPD medium, released into YPD medium containing nocodazole at 37°C to arrest cells in G2 and subjected to minichromosome IP assay. **(D)** Exponentially growing wild type (K23889) and *pds5-101* (K24030) 6C strains were arrested in G2 by addition of nocodazole at 25°C and shifted to 37°C. Samples taken at indicated times were subjected to the minichromosome IP assay. The data shown are from the same Southern blot, with irrelevant lanes removed. **(E)** Exponentially growing tetraploid cells containing 4 copies of 6C cohesin with a tag on just one of the 2C Scc1 copies (24561) and tetraploid cells containing 3 copies of 5C cohesin and one copy of 6C cohesin with a tag on just the 2C Scc1 (24560) were subjected to the minichromosome IP assay. The intensities of the CM and CD bands quantified using AIDA Image Analyzer software are plotted as % of the total lane intensities. See also Figure S1J and S1H. **(F)** Exponentially growing 6C strains containing ectopically expressed versions of 2C Scc1, K24205 (wild type) and K26413 (V137K) were arrested in G2 with nocodazole and then subjected to the minichromosome IP assay. **(G)** 6C Strains with the temperature sensitive *scc1-73* allele at the endogenous locus and either wild type (2C) Scc1 (K26600) or (2C) scc1 V137K mutant (K26591) at an ectopic locus were arrested in G1 at 25°C in YPD medium, released into YPD medium containing nocodazole at 37°C to arrest cells in G2 and subjected to the minichromosome IP assay. See also Figure S1I.

Importantly, *wpl1Δ eco1Δ* cells also formed both CMs and CDs efficiently (Fig. 2B), proving that releasing activity is responsible for the absence of CDs in cells lacking Eco1 and demonstrating that Wapl is not required for cohesin loading. Assuming that suppression of *eco1Δ* lethality by *wpl1Δ* stems at least partly from restoration of cohesion, then this result demonstrates a striking correlation between CD formation and cohesion.

While essential for maintaining cohesion, Pds5 is not required for loading (Panizza et al., 2000; Toth et al., 1999). Accordingly, *pds5-101* cells that underwent DNA replication at 37°C form CMs but no CDs (Fig. 2C). Interestingly, CM accumulation was marginally greater in *pds5-101* cells possibly because of the abrogation of releasing activity by Pds5 inactivation (Fig. 2C). It has been suggested that loss of cohesion caused by inactivation of Pds5 during G2/M despite persistence of cohesin on chromosomes is evidence that cohesin cannot hold sister DNAs together by entrapping them within a single cohesin ring (Tong and Skibbens, 2014). According to this point of view, cohesion is mediated by the interaction of two separate rings that individually entrap each sister DNA with Pds5 mediating the dimerization of cohesin rings (handcuff model). Inactivation of Pds5 is then postulated to destroy this interaction. If this were true and CD formation reflected an aspect of cohesin function unrelated to cohesion, then inactivation of Pds5 in postreplicative cells should not necessarily affect CD formation. To address this, we arrested 6C *pds5-101* cells in G2/M at the permissive temperature and then shifted the cells to the restrictive temperature, which is known to destroy cohesion (Panizza et al., 2000). This led to a reduction of CDs but not CMs (Fig. 2D). Thus, loss of cohesion during G2/M in *pds5* mutants is accompanied by loss of CDs, extending yet again the correlation between cohesion and CDs.

To address in a more definitive fashion whether individual hetero-trimeric rings or versions containing two or more kleisin subunits hold CDs together, we created two tetraploid strains that either contain four copies of covalently circularisable cohesin (4×6C) or three copies of 5C and one copy of 6C cohesin. If individual cohesin rings held CDs together, the ratio of CDs to CMs should be unaltered by any reduction in the fraction of circularisable cohesin. However, if CDs were mediated by oligomeric cohesin containing two (or more) Scc1 subunits then the fraction should be one quarter (or less) in the case of 6C/5C/5C/5C tetraploids, compared to 4×6C controls. In fact, the ratios in these two strains were very similar (Fig. 2E and S1J). The same was true in diploid cells analyzed in a similar fashion (Fig. S1H), implying that rings containing only a single copy of Scc1 to hold the two DNAs within CDs together.

### Cohesin rings collaborate to form CDs

In the course of exploring more systematically the relationship between CDs and cohesion, we analyzed cells expressing an *scc1* mutant *(V137K)* that is defective in binding Pds5 (Chan et al., 2013). To do this, we constructed strains expressing a single version of Smc1 and Smc3, each containing a pair of cysteine substitutions, and two versions of Scc1, an untagged endogenous version and an ectopic tagged version containing A547C whose N-terminal and C-terminal domains can be crosslinked to Smc3 and Smc1, respectively. In one strain, the tagged *SCC1 A547C* gene was wild type while in the other it contained *V137K.*

Due to its inability to recruit Pds5, we expected that cohesin containing scc1V137K would be able to form CMs but not CDs. To our surprise, *scc1 V137K* caused only a slight if any reduction in CDs (Fig. 2F). We considered two explanations for this surprising result. Either CD formation is insufficient for cohesion, or wild type cohesin, which cannot actually be part of the CDs associated with V137K cohesin, facilitates formation of CDs by the mutant complexes. To test the latter possibility, we replaced the endogenous wild type *SCC1* with the temperature sensitive *scc1-73* allele and measured whether V137K is still capable of forming CDs when cells underwent S phase at the restrictive temperature. Under these conditions, *scc1 V137K* supported CM but not CD formation (Fig. 2G). We conclude that wild type cohesin helps CD formation or maintenance by V137K cohesin.

Complementation between mutant *scc1* alleles had previously indicated that different cohesin complexes might interact functionally (Eng et al., 2015) and this observation was cited as evidence that cohesion is instead mediated by a pair of cohesin complexes. Our demonstration that wild type Scc1 enables Scc1V137K to form CDs provides an alternate interpretation, namely that two or more cohesin rings collaborate to produce cohesive structures that contain sister DNAs held within individual rings. This does not exclude the possibility that multiple cohesin rings collaborate through lateral interactions to form CDs (Eng et al., 2015).

### Efficient entrapment of DNAs when Smc and kleisin subunits are fused

If co-entrapment of sister DNAs mediates sister chromatid cohesion, then the cohesin ring must somehow open up, creating a gate through which DNAs can enter. A recent study suggested that an entry gate is created by transient dissociation of the Smc3/Scc1 interface (Murayama and Uhlmann, 2015). A problem with this claim is that it appears inconsistent with the fact that Smc3-Scc1 fusion proteins are functional in yeast (Gruber et al., 2006). Scc1-Smc1 fusions are also functional, casting doubt on either Smc-kleisin interface being an obligatory entry gate.

It is known that Smc3-Scc1 fusion proteins are capable of forming CDs (Haering et al., 2008). To address their efficiency, we compared the ability of wild type 6C complexes to form CMs and CDs with that of 4C complexes containing an Smc3-Scc1 fusion protein with cysteine pairs at the hinge and Scc1/Smc1 interfaces but not at the Smc3/Scc1 interface. Smc3-Scc1 4C containing complexes were capable of forming CMs and CDs with a similar efficiency to that of wild type 6C complexes (Fig. 3A). Calibrated ChIP-seq showed that Smc3-Scc1 fusion proteins translocate into peri-centric sequences following loading at core centromeres (Fig. 3B), albeit less efficiently than wild type.

**Figure 3.**
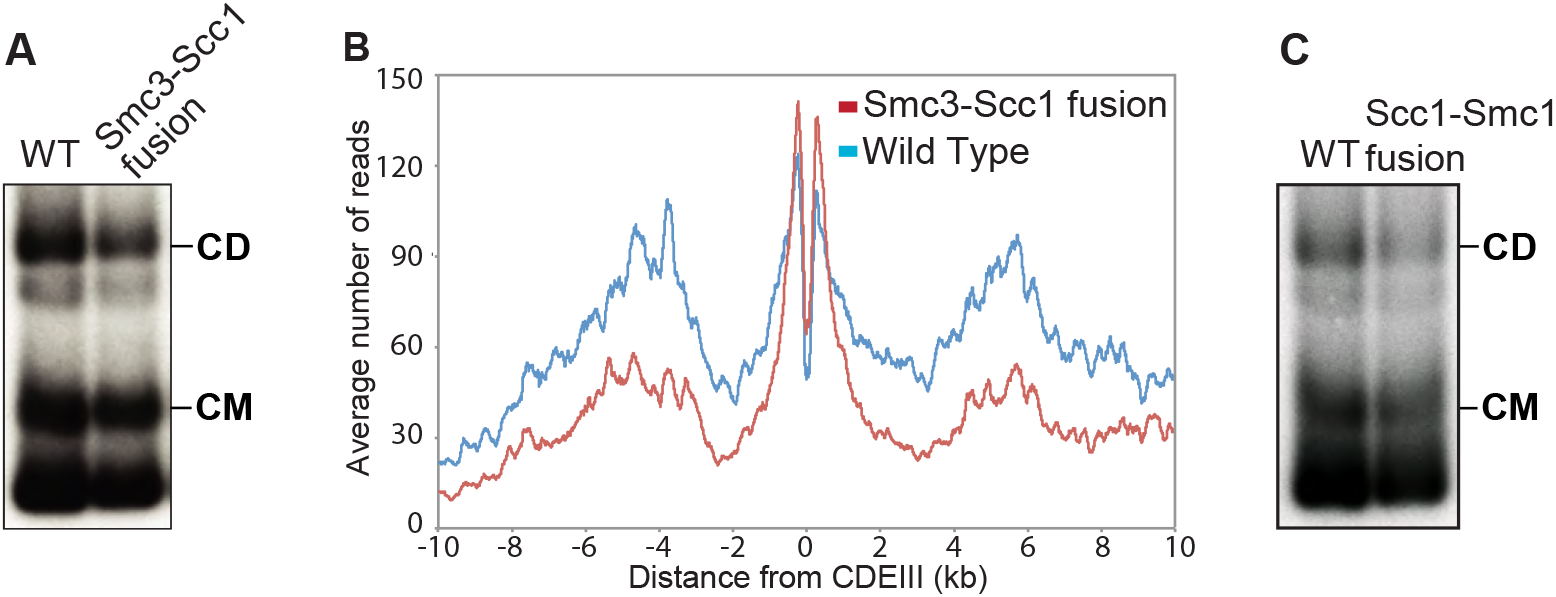
Efficient entrapment of DNAs when Smc and kleisin subunits are fused together. **(A)** A 6C wild type (K23889) strain and a strain containing 2C Smc1 and expressing an Smc3-Scc1 fusion containing cysteines in the Smc3 hinge and the Scc1 C-terminus (K24838) as the sole source of Smc1 and Scc1 were grown to log phase and subjected to the minichromosome IP assay. **(B)** Calibrated ChIP-seq of exponentially growing wild type (K23889) and Smc3-Scc1 fusion strain (K24838). ChIP profiles show the number of reads at each base pair away from the CDEIII element averaged over all 16 chromosomes. **(C)** Exponentially growing 6C wild type (K23889) strain and a 4C strain expressing a PK3-Scc1-Smc1 fusion as the sole source of Scc1 and Smc1 (K25696) were subjected to the minichromosome IP assay.

Because neither CMs nor CDs would be possible in 4C Smc3-Scc1 cells if the linker connecting Smc3 and Scc1 were cleaved, these results confirm that DNAs can enter cohesin rings via a gate other than the Smc3/Scc1 interface. They do not, however, exclude the possibility that loading can or indeed does take place via this interface. We obtained similar results with the 4C Scc1-Smc1 strains, proving that DNAs can also enter rings without opening the Scc1/Smc1 interface (Fig. 3C).

### Covalent closure of cohesin’s hinge interface fails to block loading in *Xenopus* extracts

Having established that neither Smc/kleisin interface is obligatory for cohesin loading or DNA entrapment in yeast, we addressed whether hinge opening is instead required. Our goal was to investigate the effect of crosslinking Smc1 and Smc3 moieties of the hinge using the bi-functional thiol specific reagent bBBr. Because the latter is lethal to yeast, we studied loading onto chromatin in vitro of a *Xenopus* Smc1/Smc3/Scc1/Scc3 (SA1) tetramer purified from insect cells (Fig. 4A). To distinguish exogenous and endogenous complexes, the former’s Smc3 subunit was tagged at its C-terminus with Halo (Smc3-Halo) while its Scc1 subunit contained three tandem TEV protease cleavage sites.

**Figure 4.**
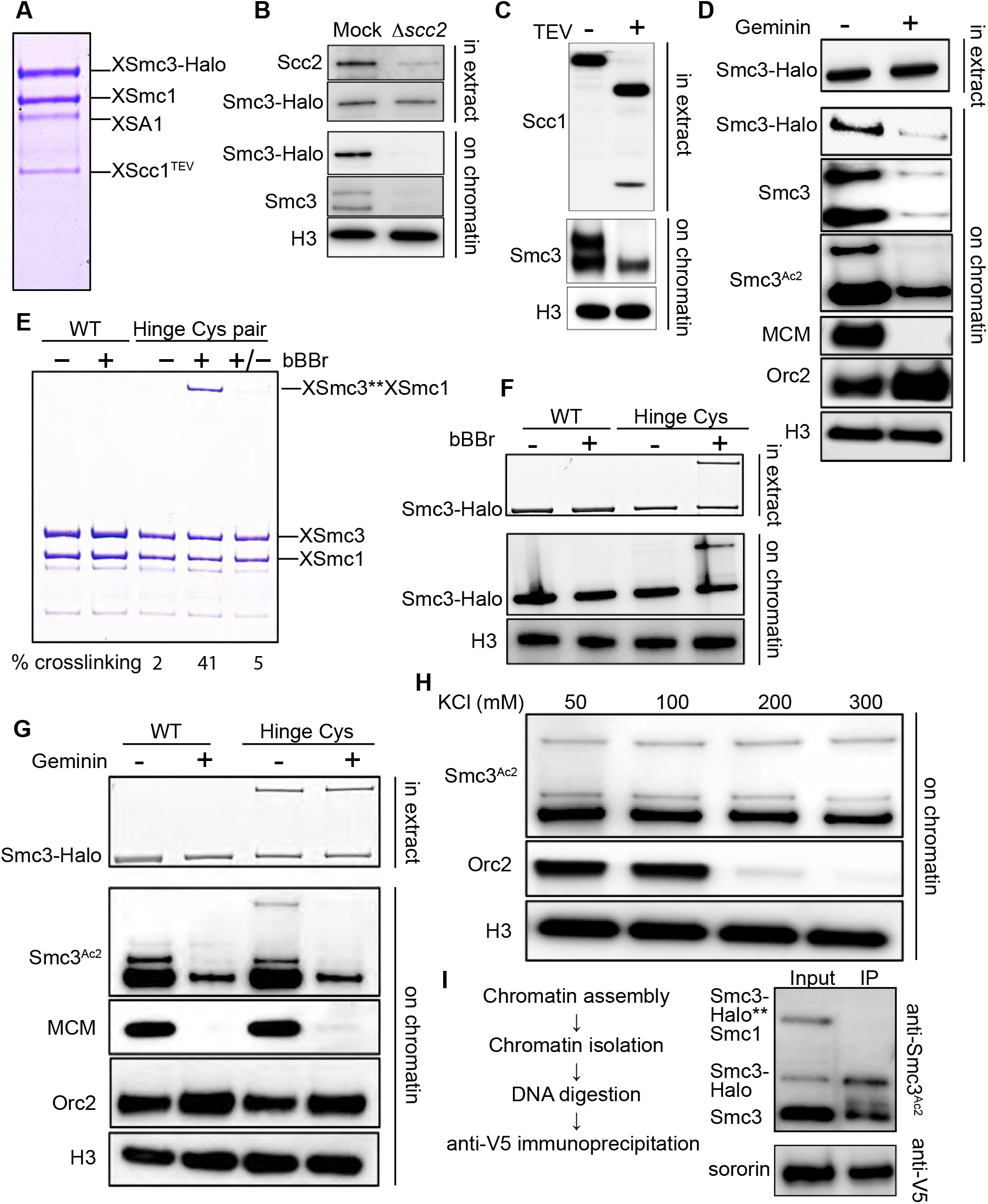
Covalent closure of cohesin’s hinge interfaces fails to block loading. **(A)** Coomassie-stained gel showing the *Xenopus* cohesin tetramer purified from baculovirus-infected *Sf-9* cells. **(B)** Mock and Scc2 depleted interphase low speed supernatant (LSS) *Xenopus* egg extracts were supplemented with the recombinant *Xenopus* tetramer, sperm nuclei and incubated at 23°C for 90 min. The isolated chromatin fraction and the soluble extracts were analyzed by Western blotting using indicated antibodies. **(C)** Recombinant *Xenopus* tetramer was treated with TEV protease or buffer for 60 min at 16°C. The reaction was then mixed with LSS interphase *Xenopus* egg extracts and treated like in (B), the chromatin and soluble fractions were analyzed by Western blotting using indicated antibodies. **(D)** LSS Interphase extract was treated with either purified recombinant geminin (60nM) or buffer for 15 min on ice. The extracts were then supplemented with recombinant *Xenopus* tetramer and sperm chromatin and treated like in (A). The chromatin and soluble fractions were analyzed by Western blotting using indicated antibodies. **(E)** Recombinant wild type cohesin (WT) and cohesin complex containing cysteine substitutions in the hinge domain (Hinge Cys) were treated with DMSO (-), 125 μM bBBr (+), or 125 μM bBBr with 10 mM DTT (+/-). Samples were also supplemented with TMR HaloTag ligand and incubated on ice for 10 min, then run on a 3-10% gradient gel. The crosslinking efficiency was quantified via TMR fluorescence. **(F)** Wild type and hinge substituted *Xenopus* tetramer was treated with DMSO or bBBr on ice for 10 min, excess crosslinker was then quenched by adding 10 mM DTT. The reactions were then supplemented with interphase extracts, TMR ligand and sperm chromatin and treated as in (B). The soluble and chromatin fractions were analysed by TMR fluorescence and indicated antibodies. **(G)** Crosslinking reactions described in (F) were supplemented with extracts pre-treated with buffer or recombinant geminin and Western blots performed as described in (D). **(H)** Hinge substituted cohesin was crosslinked and supplemented with interphase extracts and 3 ng BAC DNA/μl. After a 90 min incubation chromatin fractions were isolated, the chromatin pellets washed with buffer containing indicated amounts of KCl and analysed by Western blotting. **(I)** Hinge substituted *Xenopus* tetramer was loaded onto chromatin as in (F). The isolated chromatin pellet washed with buffer containing 300 mM KCl. The pellet was then resuspended in XB buffer supplemented with anti-V5 antibody and benzonase (1u/μl) and incubated at 12°C overnight. The immunoprecipitates were analyzed by Western blot. See also figure S7.

Addition of sperm chromatin to LSS interphase egg extracts leads to chromatin assembly, cohesin loading, acetylation of Smc3, and eventually DNA replication (Lafont et al., 2010; Takahashi et al., 2004). Like endogenous Smc3, Smc3-Halo associated with chromatin in a manner dependent on Scc2 (Fig. 4B) and was abolished by TEV-induced Scc1 cleavage (Fig. 4C). Importantly, inhibition of prereplication complex assembly by geminin addition greatly reduced association with chromatin as well as acetylation of both versions of Smc3 (Fig. 4D) (Takahashi et al., 2008).

To address whether this loading requires opening of the hinge, we produced a version of the complex containing Smc1D566C and Smc3R626C, whose cysteines at the hinge interface can be crosslinked with 40% efficiency using bBBr (Fig. 4E) (Haering et al., 2008). After crosslinking, the reaction was quenched with DTT and the purified complexes added to extracts. Importantly, the crosslinking reaction, which will modify all surface cysteines, did not adversely affect loading of exogenous wild type complex. Strikingly, Smc1D566C/Smc3R626C complexes whose hinges had been crosslinked were loaded onto chromatin (Fig. 4F) and acetylated (Fig. 4G) with similar efficiency to that of uncrosslinked complexes. Furthermore, the crosslinked and acetylated complexes were resistant to 0.3 M KCl, which abolishes most of the Orc2-DNA interaction (Fig. 4H). We conclude that complexes whose hinges cannot open are capable of loading onto chromatin in *Xenopus* extracts in a manner that permits their subsequent acetylation by Esco2.

We next asked whether chromatin bound crosslinked complexes were bound to sororin, which associates with acetylated cohesin during S phase and counteracts Wapl activity, thereby maintaining cohesion (Lafont et al., 2010; Nishiyama et al., 2010). We found that sororin’s association with chromatin becomes salt-resistant following replication (Fig. S7). We therefore assembled chromatin using bacterial artificial chromosomes (BACs), which can replicate in egg extract (Aze et al., 2016), and loaded the hinge crosslinked cohesin in interphase extracts supplemented with V5-tagged recombinant sororin. Chromatin pellets were then isolated, subjected to a salt wash and digested by benzonase. Sororin IP followed by detection of the associated acetylated Smc3 revealed that while endogenous Smc3 and the uncrosslinked recombinant Smc3 were associated with sororin, hinge crosslinked complexes were not (Fig. 4I). If salt resistant sororin binding reflects cohesion, then hinge cross-linking would appear to abrogate cohesion establishment.

### DNA entrapment is necessary for cohesion but not for loading or translocation

The finding that loading onto and translocation along chromosomes correlated with CM formation in yeast raises the possibility that both loading and translocation require entrapment of individual DNAs inside cohesin rings. If so, the ring must have an entry gate. If this does not reside at either Smc-kleisin interface, then it must reside at the hinge. And yet, our work with *Xenopus* complexes showed that they load onto chromatin even when the hinge is prevented from opening by crosslinking Smc1 and Smc3.

In fact, the results we obtained in *Xenopus* extracts with recombinant tetramers with crosslinked hinges are reminiscent of the phenotype of a version of yeast cohesin in which the positively charged central channel of its hinge domain had been neutralized by the *smc1K554D K661D smc3R665A R668A R669A* quintuple mutant *(smc1DDsmc3AAA)* (Fig. 5A). *smc1DDsmc3AAA* cohesin loads onto and translocates along chromosomes in a similar manner to wild type but it fails to generate sister chromatid cohesion and is only poorly acetylated by Eco1 during S phase (Kurze et al., 2011). If loading and cohesion are synonymous with entrapment and co-entrapment respectively, then *smc1DDsmc3AAA* cohesin should form CMs but not CDs. As expected, a 6C version of *smc1DDsmc3AAA* failed to produce CDs. More surprising, it also largely failed to form CMs (Fig. 5B and S2F) despite forming tripartite rings (Fig. S2A). Crucially, the level of minichromsome DNAs in *smc1DDsmc3AAA* immunoprecipitates was similar to wild type, showing that the mutant protein associates stably with chromatin, even though it does not entrap it (Fig. 5B). Calibrated ChIP-seq confirmed this result (Fig. S2C). To address whether *smc1DDsmc3AAA* cohesin loads and translocates normally even in the absence of wild type Smc subunits, we used calibrated ChiP-seq to measure loading of *smc1DDsmc3AAA* cohesin after cells had undergone S phase in the absence of any wild type complexes that could have aided loaded of mutant complexes. This revealed even greater amounts of mutant complexes than wild type throughout the genome. It also showed that mutant complexes loaded at *CENs* translocate like wild type into neighboring sequences (Fig. 5C).

**Figure 5.**
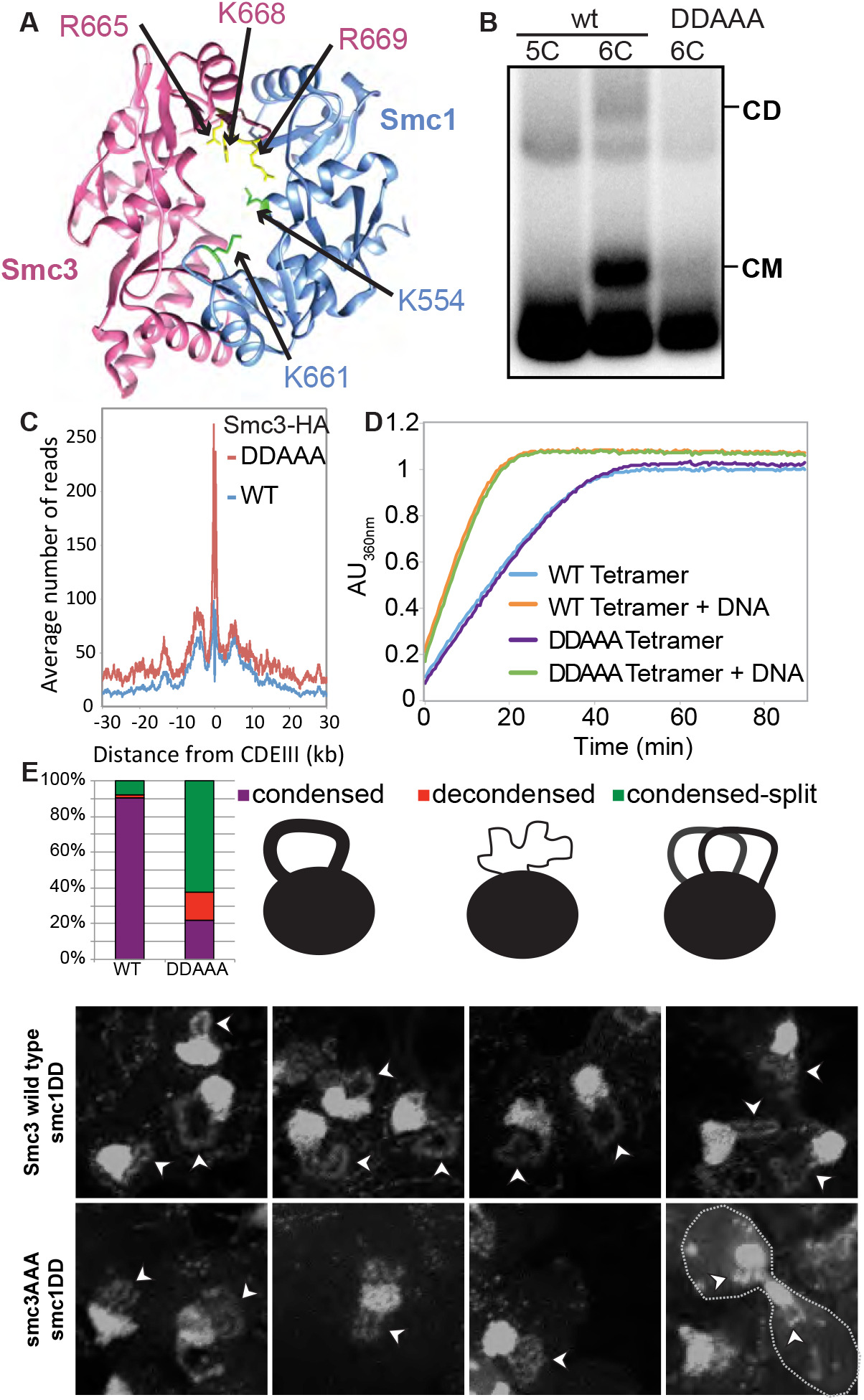
DNA entrapment is necessary for cohesion but not for loading or translocation. **(A)** Structure of the mouse hinge domain, highlighting the positively charged residues in its central channel that have been neutralized by the *smc1K554D K661D smc3R665A K668A R669A* mutations (DDAAA). **(B)** Minichromosome IP assay of exponentially growing K23644 (5C) and two 6C strains [K26210, containing a copy of wild type 2C Smc1 expressed from an ectopic locus, and K26215, containing 2C smc3AAA in the endogenous locus and 2C smc1DD in an ectopic locus (DDAAA)]. See also Figure S2A. **(C)** Strains K26797 (containing 3XminiAID-tagged smc3 in the endogenous locus and Smc3 wild type expressed from an ectopic locus), K26611 (containing 3XminiAID tagged-smc3 and smc1DD in their respective endogenous locus, and smc3AAA mutant expressed from an ectopic locus) were arrested in G1, synthetic auxin (indole-3-acetic acid) was added to 1 mM 30 min before release. The cultures were released into YPD containing 1mM auxin and nocodazole to arrest the cultures in G2 and were analyzed by calibrated ChIP-sequencing. See also Figure S2C. **(D)** ATPase activity of purified wild type and DDAAA mutant tetramer stimulated by Scc2. The rate of ATP hydrolysis was measured either in the presence or absence of DNA. **(E)** Wild type (K26797), DDAAA mutant (K26611) and K26767 (containing 3XminiAID-tagged smc3 in the endogenous locus and no additional copy of Smc3) strains were grown as in (C). 60 min after release from the G1 arrest, cultures were harvested and chromosomes spread prepared (see STAR methods). Micrographs of chromosome masses of the two strains were quantified and categorized as condensed (showing fully condensed rDNA loops), condensed-split (showing fully condensed rDNA loops that are split because of loss of cohesion) and decondensed (showing unstructured ‘puffed’ rDNA morphology). See also Figure S4.

Four important conclusions can be drawn from the behavior of *smc1DDsmc3AAA* cohesin. First, *smc1DDsmc3AAA* causes a highly specific defect in entrapping DNA. Remarkably, it causes this defect without affecting loading or translocation. Second, the Smc1/3 hinge must be intimately involved in the entrapment process, possibly acting as a DNA entry gate. Third, because the entrapment defect is accompanied by a failure to build sister chromatid cohesion, entrapment must be necessary for cohesion, which is consistent with the notion that cohesion is actually mediated by co-entrapment. Last but not least, cohesin can load onto, translocate along, and remain stably associated with chromatin in the absence of DNA entrapment. That cohesin can associate with chromatin using a non-topological mechanism as well as a topological one is a radical departure from all previous hypotheses.

### Organization of DNA into chromatid-like threads does not require entrapment of DNA by cohesin rings

The failure of *smc1DDsmc3AAA* cohesin to form CMs implies that it cannot entrap individual DNAs. However, this does not exclude the possibility that the mutant complexes associate with chromatin using a pseudo-topological mechanism, namely by entrapping loops of DNA inside cohesin rings (Fig. 7A). If so, cleavage of their kleisin subunit should release the mutant complexes from their embrace of DNA. This is indeed the case, because separase removes GFP-tagged *smc1DD smc3AAA* cohesin complexes from chromosomes during anaphase, as occurs with wild type cohesin (Fig. S2B). Given this result, we next addressed whether *smc1DD smc3AAA* cohesin is still active in organizing chromosome topology. The tandem array of rDNA repeats assemble into threads during M phase, albeit ones that are much thinner than those of conventional mitotic chromosomes (Flemming, 1880). Importantly, formation of these threads depends on cohesin (Guacci et al., 1994). Because thread formation is not dependent on sister chromatid cohesion in mammalian cells, we reasoned that the same might be true of mitotic rDNA threads in yeast. If so, and if *smc1DD smc3AAA* cohesin still possesses this thread-making activity, then post-replicative *smc1DD smc3AAA* cells should contain not one but two rDNA threads.

**Figure 6.**
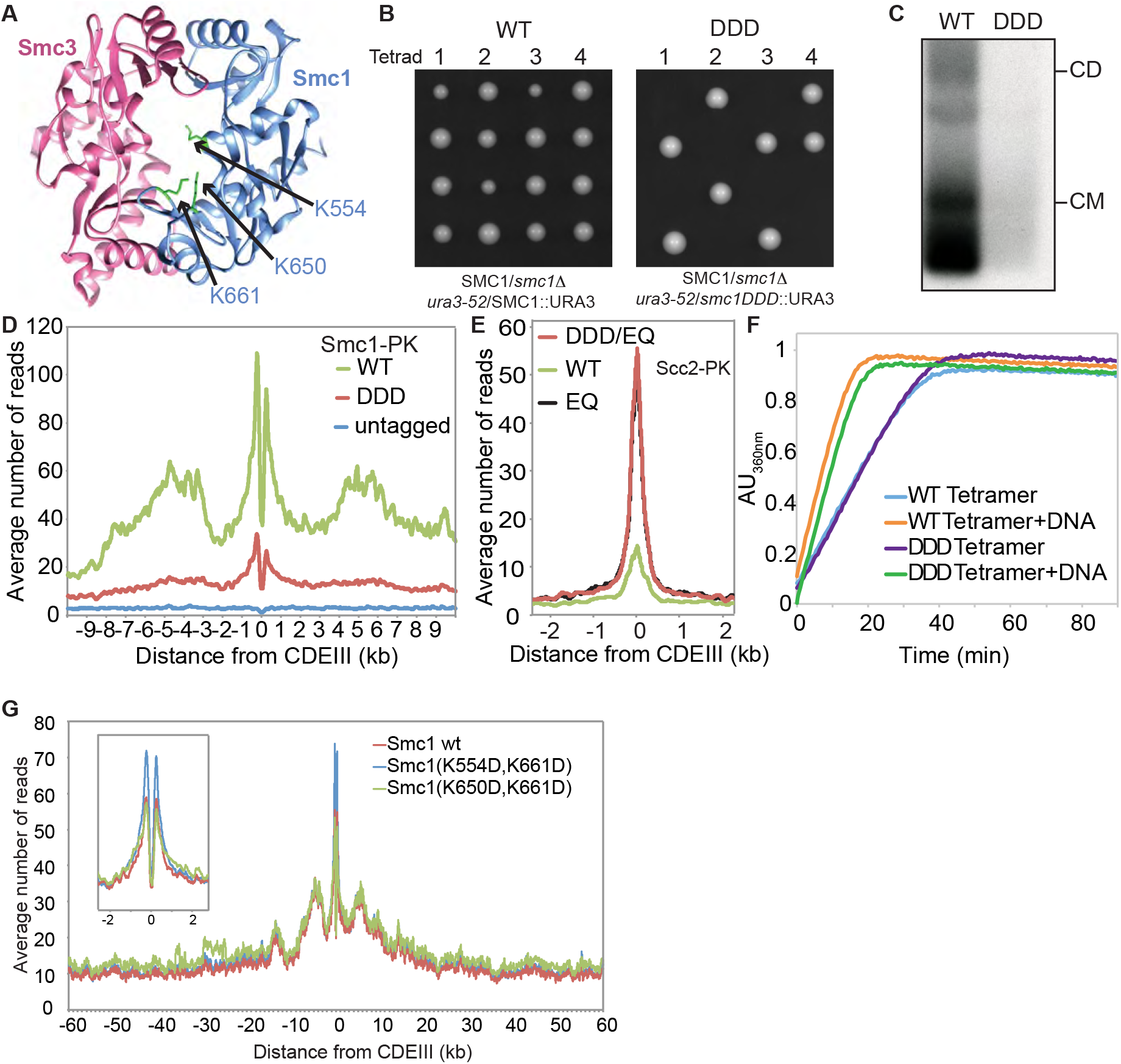
Residues within the hinge domain dictate cohesin’s ability to load and entrap DNA. **(A)** Structure of the mouse hinge depicting the corresponding *S. cerevisiae* residues within Smc1 that were mutated *(smc1 K554D K650D K661D)* (DDD). **(B)** Diploid strains *(SMC1/smc1Δ ura3-52/SMC1)* and *(SMC1/smc1Δ ura3-52/smc1DDD)* were sporulated and tetrads dissected, haploid segregants are shown. **(C)** Minichromosome IP assay of exponentially growing strains K24327 (expressing wild type 2C Smc1 from an ectopic locus) and K26610 (expressing 2C Smc1DDD mutant from an ectopic locus). **(D)** Calibrated ChIP-seq of exponentially growing strains K24327 (ectopic Smc1), K26756 (ectopic Smc1DDD) and K699 (untagged control). **(E)** Calibrated ChIP-seq of exponentially growing strains each containing Scc2-PK6, K26839 (ectopic Smc1), K26840 (ectopic smc1DDD E1158Q) and K25646 (ectopic smc1 E1158Q). **(F)** ATPase activity of purified wild type and DDD mutant tetramer stimulated by Scc2. The rate of ATP hydrolysis was measured either in the presence or absence of DNA. **(G)** Calibrated ChIP-seq of exponentially growing strains that contain a deletion of the endogenous *SMC1* gene and expressing the wild type Smc1 (K15324), smc1 K554D,K661D (K15322) or smc1 K650D,K661D mutant (K15226) from an ectopic locus.

**Figure 7.**
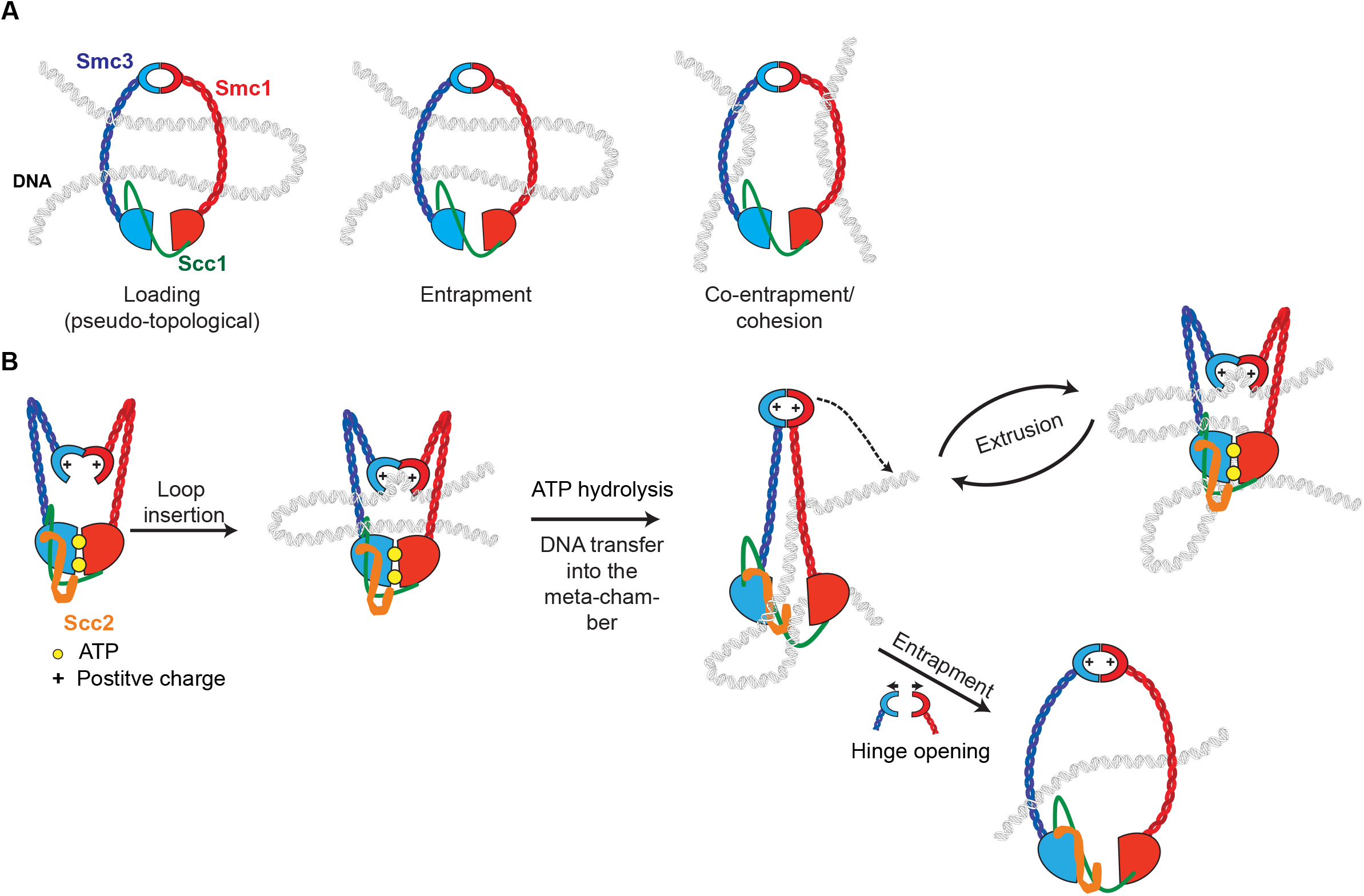
Sister Chromatid Cohesion is mediated by co-entrapment of sister DNAs inside individual rings. **(A)** Cohesin that associates with chromatin without forming cohesion can do so in two ways: one involving strict “topological” entrapment of individual chromatin fibers within cohesin rings and another that does not. We suggest that the non-topological mode involves entrapment of DNA loops within cohesin rings. **(B) Model for loop extrusion**. DNA loops can be inserted into cohesin rings without opening of any of the interfaces, the inside surface of the cohesin’s hinge acts as a DNA binding pocket whose access is regulated by its ATPase. The DNA bound to the lysine residues in the hinge is transported in a manner dependent on a cycle of ATP hydrolysis to the ATPase heads to be trapped inside cohesin’s meta-chamber. Successive cycles lead to ever-larger loops of DNA trapped pseudo-topologically at this location. The hinge rarely opens fully during this process but when it does so, this creates an opportunity for DNA entrapment

To test this, cells whose *SMC3* gene had been replaced by an AID-tagged version *(smc3-AID)*, expressing *smc1DD* from the endogenous locus and *smc3AAA* from an ectopic one were allowed to undergo DNA replication in the presence of auxin. Their behavior was compared to *smc3-AID* cells with a wild type *SMC1* gene and expressing *SMC3* from an ectopic locus (the wild type control) as well as to cells lacking an ectopic *SMC3* gene (the *smc3* mutant control). As expected most wild type cells contained a single rDNA thread, which forms a distinct loop connected to but separate from the rest of chromosome XII, which is situated within an amorphous mass of chromatin containing all 15 other chromosomes, while cells that had undergone S phase without Smc3 lacked discernable rDNA loops (Fig. S4). Remarkably, cells that had undergone S phase expressing only *smc1DD smc3AAA* cohesin frequently (68%) contained a pair of thin rDNA loops (Fig. 5E and S4). This implies that *smc1DD smc3AAA* cohesin can organize individual rDNA into threads but not hold sister rDNA threads together.

If threads are formed by the process of loop extrusion (LE), as is currently believed, then it would appear that *smc1DD smc3AAA* cohesin is still capable of LE. If LE is driven by cohesin complexes that hydrolyze ATP when associated with Scc2 instead of Pds5(Petela et al., 2017), then *smc1DD smc3AAA* cohesin should remain active as an Scc2-dependent ATPase. To test this, we purified wild type *(SMC1 SMC3 SCC1 SCC3)* and mutant *(smc1DD smc3AAA SCC1 SCC3)* tetramers, and compared their ATPase activity alongside Scc2 and in the presence and absence of DNA. Crucially, *smc1DD smc3AAA* had no effect on ATPase activity in response to DNA (Fig. 5D).

### Highly conserved lysine residues inside Smc1/3 hinges are required for all types of cohesin loading

To address if cohesin’s hinge might be involved in all aspects of cohesin’s chromosome organization and not merely the DNA entrapment intrinsic to chromatid cohesion we undertook a more systematic analysis of the role of basic residues within cohesin’s hinge. Smc1K554 and K661 are in fact part of a triad of highly conserved lysine residues, including K650, residing within the Smc1 moiety of the hinge’s lumen (Fig. 6A and S3). All double mutant combinations involving lysine to aspartate substitutions, namely *smc1K554D K650D, smc1K554D K661D,* and *smc1K650D K661D* are viable. Indeed, calibrated ChIP-seq showed that neither *smc1K554D K650D* nor *smc1K650D K661D* had any appreciable effect on cohesin’s association with the genome, either around centromeres or along chromosome arms (Fig. 6G).

In contrast, the *smc1K554D K650D K661D* triple mutant *(smc1DDD)* was lethal (Fig. 6B), greatly reduced cohesin’s association with chromatin throughout the genome (Fig. 6D), and abolished co-precipitation of minichromosome DNA with cohesin as well as formation of CMs and CDs when present as a 6C version (Fig. 6C). Treatment of cells with the 6C version of *smc1DDD* showed that chemical circularization of the triple mutant was identical to wild type, showing that the triple mutation does not adversely affect Smc1/3 hinge dimerization or indeed association between Smc1 and Smc3 ATPase domains with C- and N-terminal domains of Scc1, respectively (Fig. S2A). Because *smc1DD smc3AAA* reduces the off-rate of isolated Smc1/3 hinge complexes (Kurze et al., 2011), we used a competition crosslinking assay to measure this property but found that *smc1DDD* had no effect (Fig. S2E).

Analysis of mutations like *smc1Ell58Q* and *smc3Ell55Q* that block ATP hydrolysis has revealed two steps in the loading reaction at *CENs.* The first is association with *CENs* of cohesin whose ATPase heads have engaged in the presence of ATP while its kleisin subunit binds Scc2 instead of Pds5. A subsequent step involves conversion of this unstable intermediate into a complex that translocates up to 30 kb into neighboring peri-centric sequences, while remaining stably associated with chromatin. Formation of the unstable Scc2-bound intermediate at *CENs* can be detected using calibrated ChIP-seq, by measuring enhancement by *smc1Ell58Q* of Scc2’s association with *CENs* (Petela et al., 2017). Importantly, the enhanced recruitment of Scc2 to *CENs* in *smc1DDD smc1Ell58Q* expressing cells was identical to that in *smc1Ell58Q* expressing cells. *smc1DDD* therefore affects the second and not the first step in the loading reaction. Because *smc1DDD* has no effect on ATPase activity induced by Scc2 in vitro (Fig. 6F), we conclude that *smc1DDD* does not affect the ATP hydrolysis cycle per se but instead a change in cohesin’s conformation that normally accompanies hydrolysis of ATP bound to its ATPase heads, presumably involving the hinge and possibly also the associated coiled coils. It is remarkable that mutation of any one of three conserved lysines is sufficient to reduce wild type levels of loading in double *smc1DD* mutants to lethally low levels. None of the three conserved lysines has a unique role and all make “equivalent” contributions. Positive charge per se and not precisely where it is situated within the hinge’s lumen appears to be crucial.

To address why *smc1DD smc3AAA* merely blocks entrapment while *smc1DDD* hinders all types of loading including entrapment, we created an *smc1DDD smc3AAA* sextuple mutant. Calibrated ChiP-seq revealed that *smc3AAA* cannot ameliorate *smc1DDD’s* loading defect (Fig. S2D). In other words, *smc1K650D* is epistatic to *smc3AAA* in *smc1K554D K661D* cells. If loading, translocation, and loop extrusion are functions common to condensin and cohesin, while DNA entrapment is an activity that arose subsequently, from a modification of the hinge’s role during loading/translocation/LE, then our data suggest that the “additional” cohesin-specific entrapment function is more easily abolished in *smc1DD* cells than the ancestral LE activity.

## Discussion

### Re-evaluation of the ring model

Elucidation of cohesin’s basic geometry led to the notion that sister DNAs are held together by co-entrapment inside a tripartite ring formed by pairwise interactions between Smc1, Smc3, and Scc1. This is known as the ring model. We describe here the first systematic attempt to test a key prediction of the ring model, namely that dimeric DNAs catenated by cohesin rings in this manner (CDs) should invariably be found in post-replicative cells that that have generated cohesion while individual DNAs catenated by cohesin rings (CMs) should always be formed when cohesin is known to load onto chromosomes. With the creation of a wide variety of cohesin mutations, this undertaking had become timely. This approach was both powerful and rigorous as only a single counter-example would be sufficient to disprove either hypothesis. Our results reveal a perfect correlation between formation of CDs and cohesion in vivo. Indeed, any other hypothesis would have to explain why co-entrapment not only exists but also correlates with cohesion. The case for CDs being synonymous with cohesion is strengthened by our finding that the a mutant version of cohesin that has a specific defect in all forms of entrapment, namely *smc1DD Smc3AAA,* is as predicted defective in building cohesion.

In contrast, despite a strong correlation between cohesin loading in vivo and CM formation, our approach revealed a glaring counter-example, namely cohesin complexes with multiple mutations *(smc1DD Smc3AAA)* that reduced the positive charge of the small lumen within the Smc1/3 hinge. *smc1DD Smc3AAA* cohesin loads onto and translocates along chromatin as well if not better than wild type and yet it largely fails to form CMs. This finding suggests that when cohesin associates with chromatin without forming cohesion it can do so in two ways: one involving strict “topological” entrapment of individual chromatin fibers within cohesin rings (as detected by CMs) and another that does not. It seems implausible to imagine that the non-topological association is an artifact caused uniquely by *smc1DD Smc3AAA.* The most parsimonious explanation is that wild type cohesin uses both non-topological and topological modes and that *smc1DD Smc3AAA* can perform the former but not the latter. It is important to point out that our gel electrophoresis results do not exclude the possibility that the non-topological mode involves entrapment of DNA loops instead of individual DNA segments (Fig. 7A). Because of its topological nature, we will refer to loop entrapment as “pseudo-topological”.

### Topological entrapment requires a DNA entry gate

If cohesion is mediated by co-entrapment, then the cohesin ring must transiently open up to permit DNA entry. Where then is the DNA entry gate? If we assume that there is only one, then it cannot be situated at either of the two Smc-Scc1 interfaces because, as we show here, DNAs still enter cohesin rings containing either Smc3-Scc1 or Scc1-Smc1 fusion proteins. This leaves the Smc1/3 hinge interface. Apart from a process of elimination, is there any evidence for this location? Unlike the Smc/kleisin interfaces, it is impossible to block hinge opening by making gene fusions and we therefore addressed the issue using two different types of approach. The first was genetic. In principle, it should be possible to inactivate the gate by mutating residues within it. We suggest that the simplest explanation for the phenotype of the *smc1DD Smc3AAA* hinge allele is that it prevents entrapment, either by blocking opening or passage of DNA through it. The second approach involved testing the effect of thiol-specific crosslinks across the Smc1/3 hinge interface of *Xenopus* complexes. Consistent with the *smc1DD Smc3AAA* phenotype, this had no effect on loading but blocked association between chromosomal cohesin and sororin. If sororin association is a mark of cohesive complexes, then it would appear that sealing the hinge interface is sufficient to prevent their formation. Though less rigorous, the finding that holding Smc1/3 hinge halves together using rapamycin-mediated FKBP12-FRB complexes in yeast also points in this direction (Gruber et al., 2006).

### Cohesin does not need a DNA entry gate for loading or translocation

Another important implication of the *smc1DD smc3AAA* phenotype is that the whole notion of a DNA entry gate being necessary for cohesin loading is fundamentally misconceived. If loading is not usually accompanied by DNA’s topological entrapment, then there is simply no need for an entry gate created by the transient dissociation of Smcs from each other or from Scc1. The latter is merely necessary for co-entrapment of sister DNAs that mediate sister chromatid cohesion but not for loading, translocation, or formation of chromatid-like threads and therefore possibly not for LE. If so, topological entrapment involving a DNA entry gate may be a feature unique to cohesin and may be lacking in all other Smc/kleisin complexes, including cohesin’s closest relative condensin.

### The hinge is required for loading as well as DNA entrapment

One of our most unexpected findings is that the hinge has a key role in loading cohesin onto chromosomes as well as DNA entrapment. A remarkable aspect of the loading function is its disruption through substitution by acidic residues of three highly conserved Smc1 lysine residues inside the hinge’s lumen *(smc1DDD).* Because loading is unaffected by mutating two out three residues, all three must equivalent roles, including one, namely K650, that unquestionably points inside the hinge’s lumen. Because *smc1DDD* cohesin forms normal rings in vivo and is fully active as an ATPase in vitro, we suggest that its loading defect arises because it fails to execute an action that normally accompanies the ATP hydrolysis cycle during the loading process. We cannot exclude the possibility that the lysines within the hinge lumen facilitate conformational changes in the Smc coiled coils emanating from the hinge, that take place in response to ATP binding/hydrolysis by their heads as envisaged in bacterial SMCs (Minnen et al., 2016). Because they do not affect the integrity of the hinge or formation of hetero-trimeric rings, they are more likely to act through their transient exposure to a negatively charged substrate, such as DNA. We therefore favor the notion that a change in the hinge’s conformation is required for loading and translocation as well as DNA entrapment.

An important clue as to its nature is that *smc1K554D K661D* complexes, which behave like wild type, are converted to ones that cannot load at all by *smc1K650D* but to ones that can load but not entrap by *smc3R665A R668A R669A.* The implication is that there is something in common between the hinge conformational changes necessary for loading and the process of entrapment. If the latter involves hinge opening, then the former might involve a more modest change that merely exposes Smc1K554 K650 K661 to their substrate. We speculate that being normally hidden inside the hinge’s lumen, these lysines are only exposed to their substrate (possibly DNA) transiently, at a certain stage of the ATP hydrolysis cycle mediated by Scc2. In other words, the inside surface of the cohesin’s hinge may act as a DNA binding pocket whose access is regulated by its ATPase.

### The ring model revised

In formulating a model to explain our results, we made the following assumptions. 1) Cohesin loading is invariably accompanied by translocation and involves pseudo-topological entrapment because this explains why cohesin forms a ring and why kleisin cleavage releases cohesin from chromatin. 2) Cohesin’s ability to form chromatid-like threads is driven by LE, which could be driven either by a locomotive device that walks along DNA (Terakawa et al., 2017) (Fig. S5A) or by a pump-like action (Diebold-Durand et al., 2017; Terakawa et al., 2017) (Fig. S5B), or as described below by a mechanism involving both. 3) If cohesin’s coiled coils are the legs using for walking along DNA, then its two feet are likely to be its hinge domain at one end and Hawks associated with its kleisin subunit and/or ATPase domains at the other end. 4) During its loading/translocation cycle, DNA is trapped in a pseudo-topological sense inside a meta-chamber (Diebold-Durand et al., 2017) formed by association of N- and C-terminal kleisin domains with Smc3 and Smc1 ATPase heads that have engaged in the presence of ATP. Fig. 7B describes a speculative model that takes these features into account. Briefly, we envision that the lysines in cohesin’s hinge bind to DNA, which is transported in a manner dependent on a cycle of ATP hydrolysis to Hawks associated with its kleisin subunit and its ATPase domains. To explain the processive nature of loop extrusion, we envision that DNAs transported to the ATPase heads are subsequently trapped inside cohesin’s meta-chamber and that successive cycles lead to ever-larger loops of DNA trapped pseudo-topologically at this location. The hinge rarely opens fully during this process but when it does so, this creates an opportunity for DNA entrapment, a process that must occur at high efficiency at replication forks when cohesion is established. Remarkably, *smc1DD Smc3AAA* disrupts replication-dependent acetylation of Smc3’s ATPase head (Kurze et al., 2011) as well as DNA entrapment, which is consistent with the notion that hinges transiently interact with Smc heads.

We note that the lumen within condensin’s hinge also contains highly conserved basic residues (Fig. S6). One of these corresponds to Smc1K650 (Smc4R806) while the other four are unique to condensin. A role for Smc hinges in the loading and translocation may therefore apply to all Hawk-containing Smc/kleisin complexes. This could conceivably extend to Kite-containing complexes (Palacek & Gruber, 2015). There is a striking similarity between the phenotype caused by *smc1DDD* and that by alterations in the length of Smc coiled coils in *B. subtilis* (Burmann et al., 2017). Both affect loading and translocation without adversely affecting ATPase activity in vitro or indeed association of EQ complexes with loading sites. Thus, they are both specifically “defective in coupling ATP hydrolysis to essential DNA transactions on the chromosome” (Burmann et al., 2017). It is therefore conceivable that the event that is disrupted by *smc1DDD* shares features with that disrupted by altering the phase of Smc coiled coils in *B. subtilis.* Elucidating the mechanism by which ATP binding/hydrolysis bring about conformational changes in the hinge and coiled coils has the potential to reveal the universal enzymatic principle that organizes chromosomal DNA in most organisms on this planet.

## Author Contributions

M.S., J.C.S., and K.A.N. designed and conducted experiments and wrote the manuscript. V.C. designed experiments. N.J.P., M.W., S.O., T.G.G., K-L. C., B.H., J.M., J.C., M.V., A.V.G., and A.K. conducted experiments.

## Acknowledgements

We are grateful to Katsuhiko Shirahige, Julian Blow and Rob Klose for supplying antibodies, and to Byung-Gil Lee and Jan Löwe for sharing plasmids and protocols. We thank all members of the Nasmyth group for valuable discussions and technical assistance, in particular Jean Metson and Antonio Valdés Gutiérrez. This work was funded by the Wellcome Trust (to K.N.), ERC (to K.N.) and Cancer Research UK (C573/A 12386 to K.N.). V.C is funded by the Associazione Italiana per Ricerca sul Cancro (AIRC), ERC and the Harvard-Armenise foundation.

